# Ultrasound-induced Particle Dynamics in Pathological Vascular Vortices

**DOI:** 10.1101/2025.10.23.684129

**Authors:** Mahmoud Medany, Nitesh Nama, Daniel Ahmed

## Abstract

Disturbed flow is a hallmark of diseased vasculature, yet its influence on particle behavior under external actuation remains poorly understood. We uncover distinct behaviors of microparticles under disturbed flow when exposed to ultrasound, revealing selective trapping and aggregation phenomena that differ fundamentally between soft and rigid particles. Using microfluidic models of disturbed vascular flow, we show that microbubbles become trapped at the eye of vortices and self-assemble via ultrasound-induced forces. As clusters grow to a critical size, they are ejected and adhere to the wall opposite the ultrasound source, forming nuclei that progressively occupy aneurysm cavities—a mechanism that could enable targeted, noninvasive treatment. These findings reveal an unexplored interplay between ultrasound and hydrodynamic forces, offering a new strategy for ultrasound-guided therapeutic delivery in vascular disease.

**One-Sentence Summary:** Vortices are common in diseased arteries, yet we don’t know how therapeutic carriers behave in them under ultrasound. We discover that microbubbles self-assemble in vortex cores, then eject and anchor to vessel walls—revealing a new transport regime with therapeutic potential.

## Introduction

Disturbed flow, marked by separation, recirculation, and vortex formation, naturally arises at geometric complexities such as arterial bifurcations and the Circle of Willis, and it is a hallmark of diseased vasculature, including atherosclerotic segments and aneurysm cavities^1,2^. While the fluid mechanics of these regimes have been investigated^3–5^, the behavior of particles within disturbed flows—especially under external fields^6–8^—remains largely unexplored. Understanding this regime is essential not only for advancing our understanding of the fluid physics, but also for developing new strategies in targeted drug delivery, embolization, and ultrasound-guided therapies^9–11^.

Prior work in inertial microfluidics showed that suspended particles migrate across streamlines to stable equilibria due to lift forces in confined channels^12–14^, and that geometric features such as T-junctions can passively retain particles in recirculation zones^15^. However, these investigations largely excluded external forcing and focused on symmetric or steady-state flows. External fields have since been combined with laminar microchannel flows to enable advanced particle manipulation ^16–20^, but these approaches do not capture the complex physics of disturbed flow environments, where vortices, separation zones, and oscillatory shear fundamentally reshape particle–field interactions. The combined influence of disturbed flow and external actuation therefore defines a new, unexplored regime in which particle motion is dictated by the nonlinear coupling between background flow structures and applied forces.

Here, we experimentally investigate this coupled regime by examining microparticle dynamics in disturbed flows under external acoustic actuation. Using ultrasound (US) as a model field within microfluidic systems designed to reproduce disturbed flow, that occurs in blood vessels including aneurysm-like cavities or plaque-induced constrictions, we observed strikingly different dynamics depending on particle compressibility and the presence of actuation. Solid microparticles followed streamlines or became dispersed without sustained trapping, highlighting the limitations of purely passive mechanisms. In contrast, compressible microbubbles^21^ (MBs) were drawn into recirculating vortices, where they clustered, merged, and grew until reaching a critical size. At that point, the clusters were expelled from the vortex eye and propelled across the cavity by acoustic radiation forces, ultimately anchoring to the wall opposite the US source. We studied this behavior both experimentally and through theoretical modeling, the latter capturing the initial migration and trapping of MBs within vortex cores. This sequence of clustering, expulsion, and anchoring reveals a previously unreported regime of particle transport in disturbed flow, arising from the interplay between convective transport, vortex confinement, and external field coupling. Beyond the fundamental physics, these dynamics demonstrate how the combination of disturbed flow and US can be harnessed to accumulate therapeutic carriers within aneurysm-like cavities, suggesting new strategies for embolization and drug delivery.

## Results

### Experimental Setup

To investigate the behavior of MB swarms under disturbed flow and their US-induced dynamics, we fabricated microfluidic channels that reproduce flow disturbances seen in pathological vasculature. The conceptual overview is shown in Fig. 1A, and microfluidic realizations are shown in Fig. 1B–C (plaque-narrowed vessel and aneurysm-like cavity with recirculating vortex). Channel dimensions and operating conditions were informed by blood-flow speeds reported for the internal carotid artery (∼76–82 cm s^−1^) and the middle cerebral artery (∼58 cm s^−1^). To model intracavity recirculation, we designed an aneurysm-mimicking dome connected to a side channel positioned directly beneath the cavity opening. Numerical simulations illustrating vortex formation across this geometry are provided in Fig. S1.

**Fig. 1.**
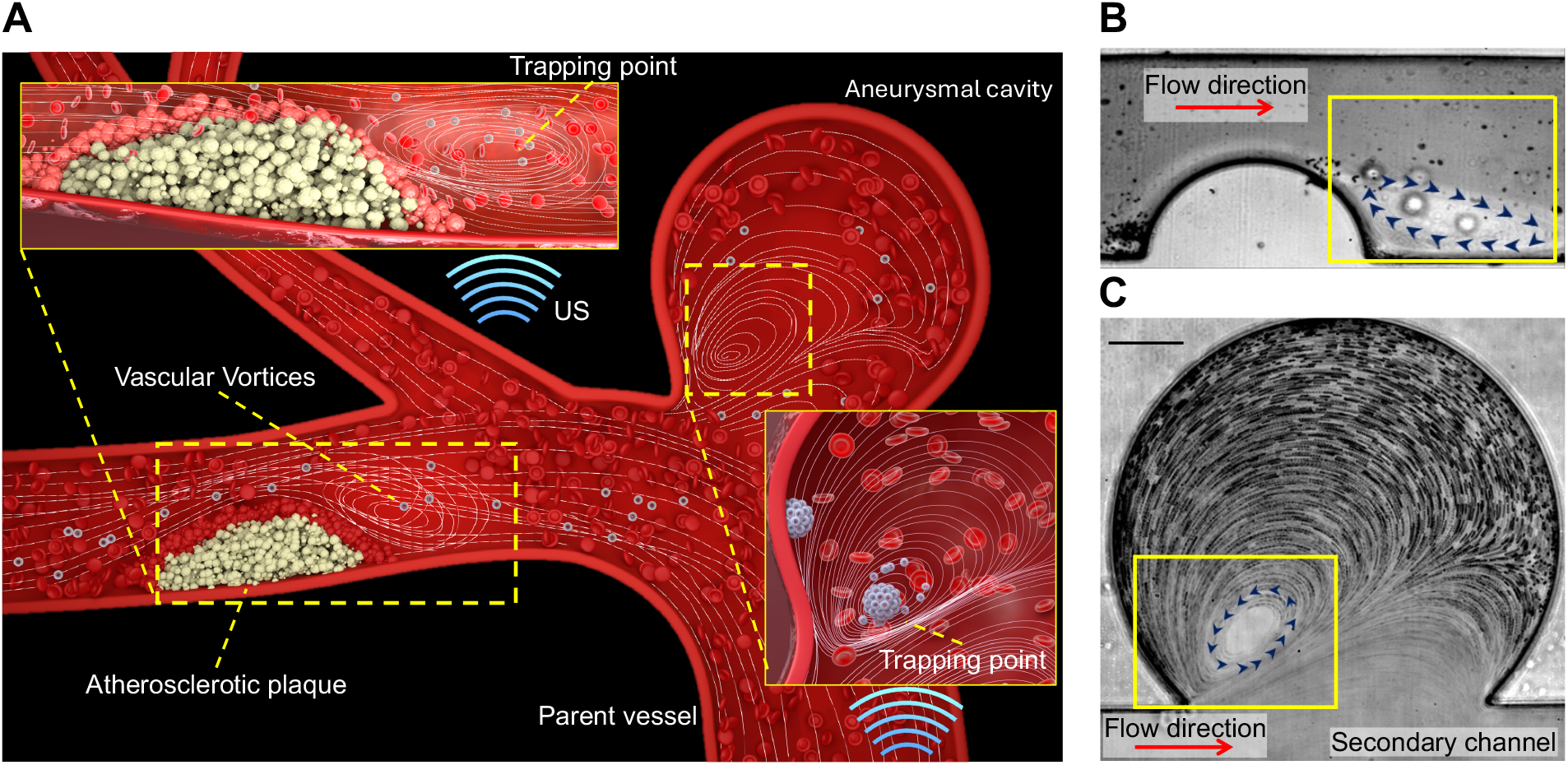
Conceptual and experimental illustration of disturbed flow in plaque- and aneurysm-representative vascular environments. **(A)** Schematic of disturbed hemodynamics in diseased vessels, showing vortices that arise downstream of atherosclerotic plaques and within aneurysmal cavities. US acts on compressible MBs within these vortical regions, concentrating them at trapping points along the low-pressure cores. **(B)** Microscope image of disturbed flow generated by a plaque-like structure inside the microfluidic model. **(C)** Microscopic view of flow patterns within an aneurysm cavity under physiological flow conditions.

A piezoelectric transducer that generated US was bonded to one sidewall of the polydimethylsiloxane (PDMS) channel, oriented orthogonally to the aneurysm cavity (Fig. S2). The transducer was driven by a function generator. To establish a predominantly traveling acoustic field and suppress wall reflections, the entire chip was immersed in a water bath. The device was mounted on an inverted microscope with high-speed imaging. Flow was imposed by a programmable pump (continuous or pulsatile), with center-line velocities from 20 to 100 cm s^−1^. For flow-field measurements, 1 µm tracer particles were used and analyzed by particle image velocimetry (PIV).

The flow profile transitioned from laminar to disturbed when the flow rate reached 30 cm s^−1^ (*Re* = *ρvL*/*μ* = 34), where Re represents the Reynolds number, *ρ* as the fluid density, *v* the velocity, L the characteristic length, and *μ* the dynamic viscosity), producing a vortex that first appeared on the left side of the dome. Increasing Re shifted the vortex centroid rightward, in agreement with the numerical simulations (Fig. S1). PIV showed a single recirculation cell: in-plane speeds peaked along the separating shear layer at the cavity mouth and decayed toward a near-stagnant eye, consistent with a Rankine-like vortex; full maps are in Fig. S2.

### Microbubble behavior under ultrasound in continuous flow

After establishing the flow regime, we investigated the behavior of MBs within the aneurysm cavity under continuous flow and US exposure. Flow rates between 60 and 80 cm s^−1^ (*Re* = 68– 100) generated a consistent microvortex within the cavity due to flow divergence at the channel– cavity junction. Under these conditions, when gas-filled MBs (2.5 µm, Sonovue) were introduced into the microfluidic chamber, they typically remained in the secondary channel (Fig.1C) and bypassed the aneurysm cavity in the absence of acoustic actuation. Upon continuous activation of US, the acoustic pressure gradient generated by the transducer pushed the MBs from the channel into the adjoining cavity. Once inside, they entered the vortex and followed elliptical trajectories, spiraling progressively toward the core. For example, at an external flow rate of 60 cm s^−1^, MB speeds ranged from ∼ 3 mm s^−1^ near the cavity entrance to ∼ 0.5 mm s^−1^ near the vortex eye; with successive cycles, speeds decreased further, culminating in entrapment at the vortex eye (Fig. 2A). High-speed measurements of individual MB velocity profiles are shown in Fig. 2B, and sequential frames illustrate their progressive convergence into the core (Fig. 2C).

**Fig. 2.**
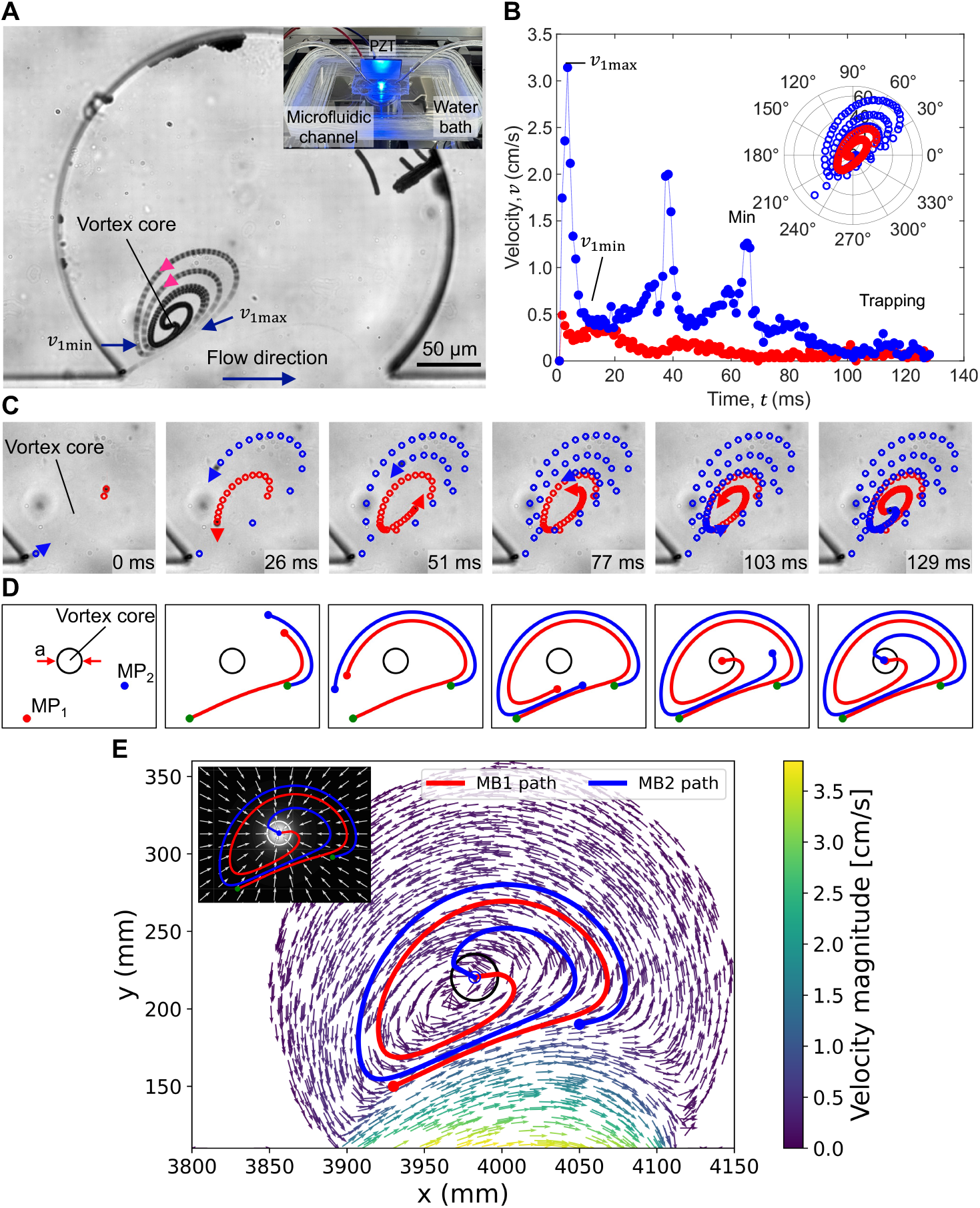
Experimental and theoretical characterization of MBs spiraling into the vortex core. (**A**) Overlay of the image sequence illustrating the MB trajectory as it approaches the trapping point. **(B**) A velocity profile with 134 points shows three peaks of velocity, while the MBs represent the high-shear flow profile, and a polar plot illustrates the radius and angle from the vortex center. (**C**) The time sequence of two MBs attracted to the vortex core as they circulate. **(D)** Simulation of two MBs released at distinct entry angles, reproducing inward spiraling and convergence to the vortex core. **(E)** COMSOL-derived flow field overlaid with simulated MB trajectories, showing robust capture across entry positions. Inset: Rankine-vortex pressure well establishing the gradient that guides MBs into the trapping region.

### Minimal model of spiraling capture, clustering, and ejection

To understand the observed spiraling capture, we consider a theoretical model that combines the measured vortex flow field with a Rankine-type pressure trap and a non-zero US forcing term. Trapping only occurs when US is active; therefore, the primary radiation force **F**_rad_ s included from the outset, and its effective magnitude grows as clusters grow (owing to its scaling with particle volume^22–24^ *V*_*c*_(*t*)). For a bubble or cluster of effective radius *R*_*c*_and volume 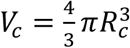 at position **x**(*t*)with velocity ***v***(*t*), the force balance is

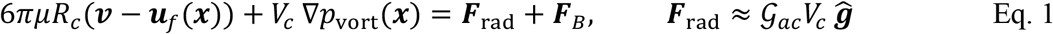

Here, *μ* is fluid’s shear viscosity; ***u***_*f*_(***x***) is the carrier flow field; and *p*_vort_(**x**)is a Rankine-like pressure surrogate with a minimum at the eye (core radius *a*, circulation Γ, spin Ω = Γ/2*πa*^2^). We define the inward confinement force **F**_*v*_ ≡ − *V*_*c*_∇*p*_vort_ where *p*_vort_ has a minimum at the eye such that **F**_*v*_ points inward. The unit vector 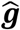points from the transducer toward the opposite wall, and 𝒢_ac_ (Pa m^−1^) encapsulates the effective acoustic contrast and the mean pressure-squared gradient. Because **F**_rad_ ∝ *V*_*c*_(*t*), the acoustic term is non-zero when US is on (driving entry and initial capture) and becomes effectively stronger as clusters grow at the eye. The additional term **F**_*B*_ captures secondary Bjerknes attractions among oscillating MBs in the core and the image-Bjerknes^25^ pull near a wall (schematics in Fig. 3A–B).

**Fig. 3.**
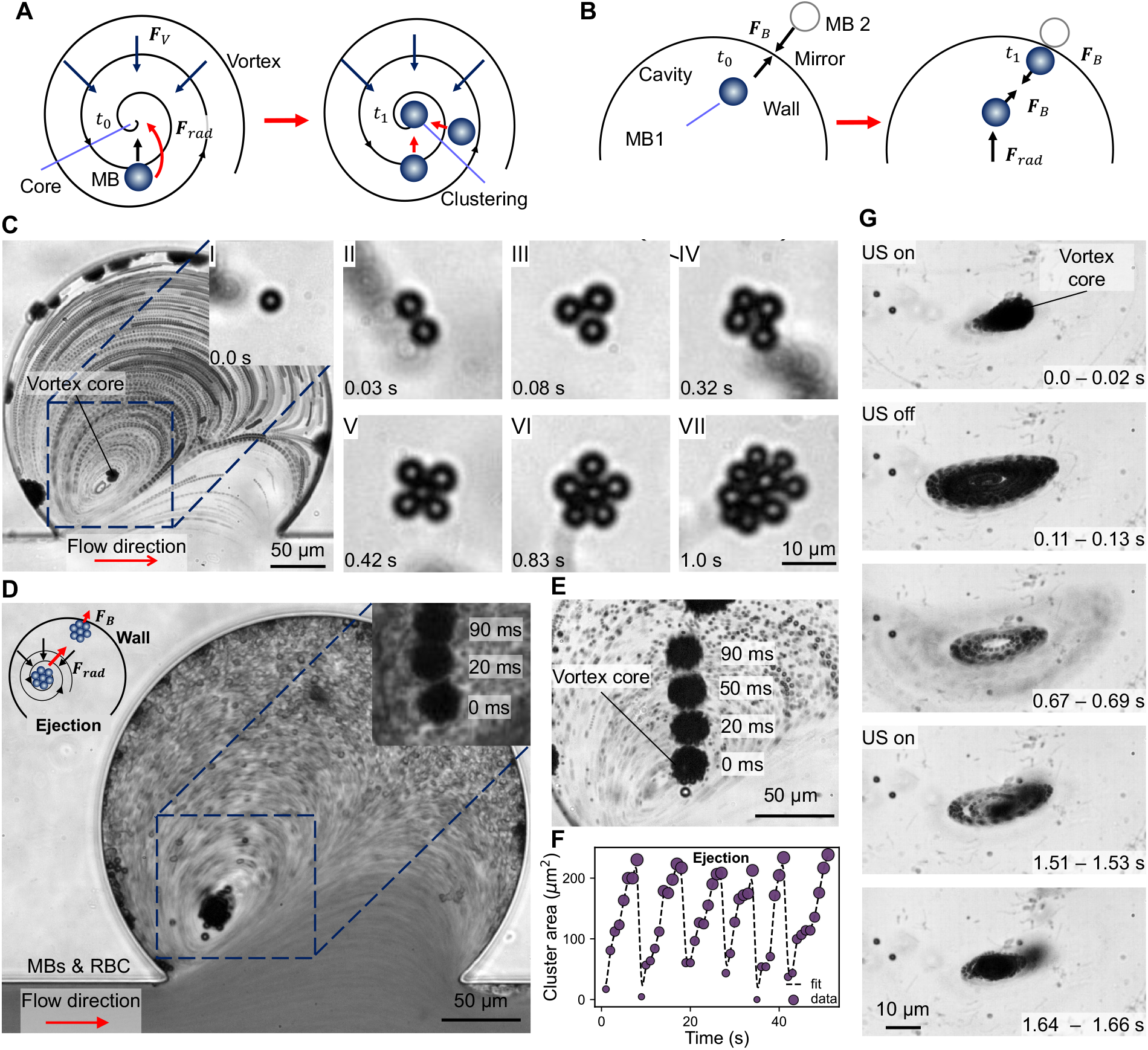
Collective dynamics of MBs under US. **(A)** Multiple MBs are drawn into the aneurysm cavity and become trapped at the vortex eye. **(B)** Schematic illustrating secondary Bjerknes forces: MB–MB attraction within the vortex core and image-mediated attraction toward the cavity wall. **(C)** Sequential frames showing progressive MB assembly at the vortex eye, evolving from doublets to triplets, quadruplets, and larger aggregates within milliseconds (scale bar = 50 µm). **(D)** Ejection of an MB cluster from the vortex core in the presence of red blood cells (RBCs), which largely follow streamlines and bypass the trap. (scale bar = 50 µm) **(E)** Time-lapse series showing that cluster ejection occurs once a critical radius (∼15 µm) is reached. (scale bar = 50 µm) **(F)** Quantification of cluster size at the moment of ejection across multiple events, showing the critical threshold for destabilization. **(G)** On/off control verifying ultrasound dependence: with US on, MBs compact and remain at the vortex eye; switching US off leads to gradual de-clustering and drift from the core; turning US on again restores trapping.

To validate the model, we simulated the motion of two MBs released at distinct entry positions within the measured flow field. Both trajectories reproduced the experimental behavior: each bubble spiraled inward along elliptical paths and converged to the vortex core (Fig. 2D). The complete COMSOL velocity field, overlaid with the simulated trajectories, shows that inward capture is robust across entry angles (Fig. 2E). The inset in Fig. 2E depicts the Rankine-vortex pressure well that establishes the gradient guiding MBs into the trapping region.

What emerged as particularly intriguing from our control experiments was the behavior of solid tracer polystyrene microparticles (1 µm) co-infused with MBs. In the absence of US, both MBs and tracer particles followed their streamlines, showing no core accumulation. Under US activation and identical conditions, however, MBs were redirected into the cavity and trapped at the vortex eye, while the trajectories of the solid microparticles remained unchanged. This difference in behavior can be attributed to particle properties: gas-filled MBs are highly compressible and oscillate under US, experiencing strong primary radiation forces and secondary-Bjerknes attractions^22–24^ that bias them toward the low-pressure eye; rigid polystyrene particles and RBCs are comparatively incompressible, respond weakly to US, and are carried by inertial drag past the eye. As MBs cluster at the eye, their effective radius increases and the acoustic bias strengthens; once a critical size is reached, clusters eject and subsequently anchor at the opposite wall via image-bubble attraction^25^. (See schematics in Fig. 3A–B.)

Once multiple MBs entered the cavity, they were consistently drawn into the vortex eye and became stably trapped. Within this low-pressure, low-shear region, US-driven interactions dominated: oscillating MBs attracted one another via secondary Bjerknes forces, MB-MB interactions initiated a progressive hierarchical assembly, with doublets, triplets, and larger aggregates forming within milliseconds (Fig. S3). With continued US, each cluster acted as a nucleus recruiting neighboring MBs from the flow, producing steady growth (∼6 MB s^− 1^; ∼30 µm^2^ s^− 1^ area increase). Under fixed conditions (18.1 V_pp_, 1.04 MHz), clusters expanded until they reached a critical radius of ∼15 µm, beyond which the trap destabilized, and the aggregate was abruptly ejected from the eye (Fig. 3C–D for assembly and ejection sequences). In experiments with red blood cells, RBCs followed streamlines and bypassed the trap, whereas MB clusters were selectively ejected once they exceeded the critical size. Ejection occurred at velocities initially as low as ∼0.25 mm s^−1^, increasing up to ∼1 mm s^−1^ just before reaching the aneurysm wall (Fig. S5), driven by the competition between drag, primary radiation forces, and secondary Bjerknes interactions, including mirror-bubble attraction to the nearby wall. Once displaced, MB clusters migrated toward the opposite wall and became stably anchored by image-mediated forces^25^.

Because the US-driven acoustic forces on an MB cluster—the primary radiation force and the secondary/image-Bjerknes term—scale with cluster volume 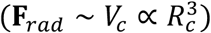 ^19-21^, whereas the viscous drag grows linearly with radius (***F***_*D*_ = 6*πμR*_*c*_ *U*), larger clusters experience an increasing outward slip relative to the background recirculation. Ejection occurs once the ejection forces (primary radiation and the image-Bjerknes force) pointing away from the eye of the vortex exceed the confining forces (hydrodynamic confinement and drag). Under our fixed driving conditions, this condition is met at a reproducible critical size (∼ 15 *μ*m; Fig. 3E–F), after which clusters depart the eye and traverse the cavity. Moreover, ultrasound dependence was reversible: switching the field off caused the compact cluster to relax and drift from the vortex eye, whereas restoring ultrasound immediately re-established trapping and re-assemble at the core (Fig. 3G).

### Progressive filling of vascular cavities by US-guided microbubbles

The ejection behavior observed in the previous experiments (Fig. 3)—where clusters released from the vortex core migrated toward and attached to the cavity wall—suggested that repeated capture– ejection cycles could gradually fill the cavity. To test this idea, we examined MBs behavior under continuous high-velocity flow. A suspension of MBs (4 µL mL^− 1^) was introduced into the secondary channel at 60 cm s^− 1^. In the absence of ultrasound, MBs followed the main streamlines and bypassed the cavity. Upon US exposure (2.03 MHz, 20 V_pp_; continuous), primary acoustic radiation forces redirected MBs into the vortex eye, where they accumulated into clusters. As the clusters grew, they reached a critical size at which the acoustic drive overcame the local vortex confinement, resulting in ejection toward the cavity sidewall. Once deposited, clusters adhered to the wall through secondary Bjerknes interactions, forming stable nuclei for further accumulation. The adhered aggregates attracted additional MBs from the circulating flow, reinforcing local trapping and promoting progressive buildup. Through successive capture, ejection, and adhesion events, the cavity became almost completely filled within approximately 15 s (Supplementary Fig. S5). Flow rate and MB concentration had a strong influence on this process. At higher flow velocities (68, 77, and 92 cm s^− 1^), the filling time was modestly extended, reaching up to 25 s at 92 cm s^− 1^ (Supplementary note 5).

To explore a more physiologically relevant disturbed-flow condition, we imposed pulsatile flow (60 bpm; 30–80 cm s^− 1^; Fig. 4A). This profile produced an oscillatory vortex within the cavity whose core shifted laterally during each beat (Fig. 4B–C). When continuous sinusoidal US waveform was applied at 1.7 MHz and 15 V_pp_, MB clusters were captured at the instantaneous vortex eye and moved synchronously with the shifting vortex core throughout the pulsatile cycle (Fig. 4D). This behavior demonstrates that the acoustic field can maintain stable trapping even as the vortex structure oscillates in both space and time. To assess whether this dynamic trapping could lead to complete cavity filling, we increased the MB concentration to match the continuous-flow experiments (4 µL mL^− 1^) and applied continuous US at 2.0 MHz and 20 V_pp_. Under these conditions, the familiar capture–growth–ejection–adhesion sequence was observed: clusters formed at the moving vortex eye, grew by recruitment, and were ejected toward the wall, where they adhered and served as nucleation sites for further accumulation. Repeated over successive pulsatile cycles, this process progressively coated the inner wall, resulting in uniform cavity filling within approximately 75 s (Fig. 4E–F). Quantification of this process was obtained from binarized image sequences, where the fraction of white pixels within the aneurysm region represented the percentage of filling.

**Fig. 4.**
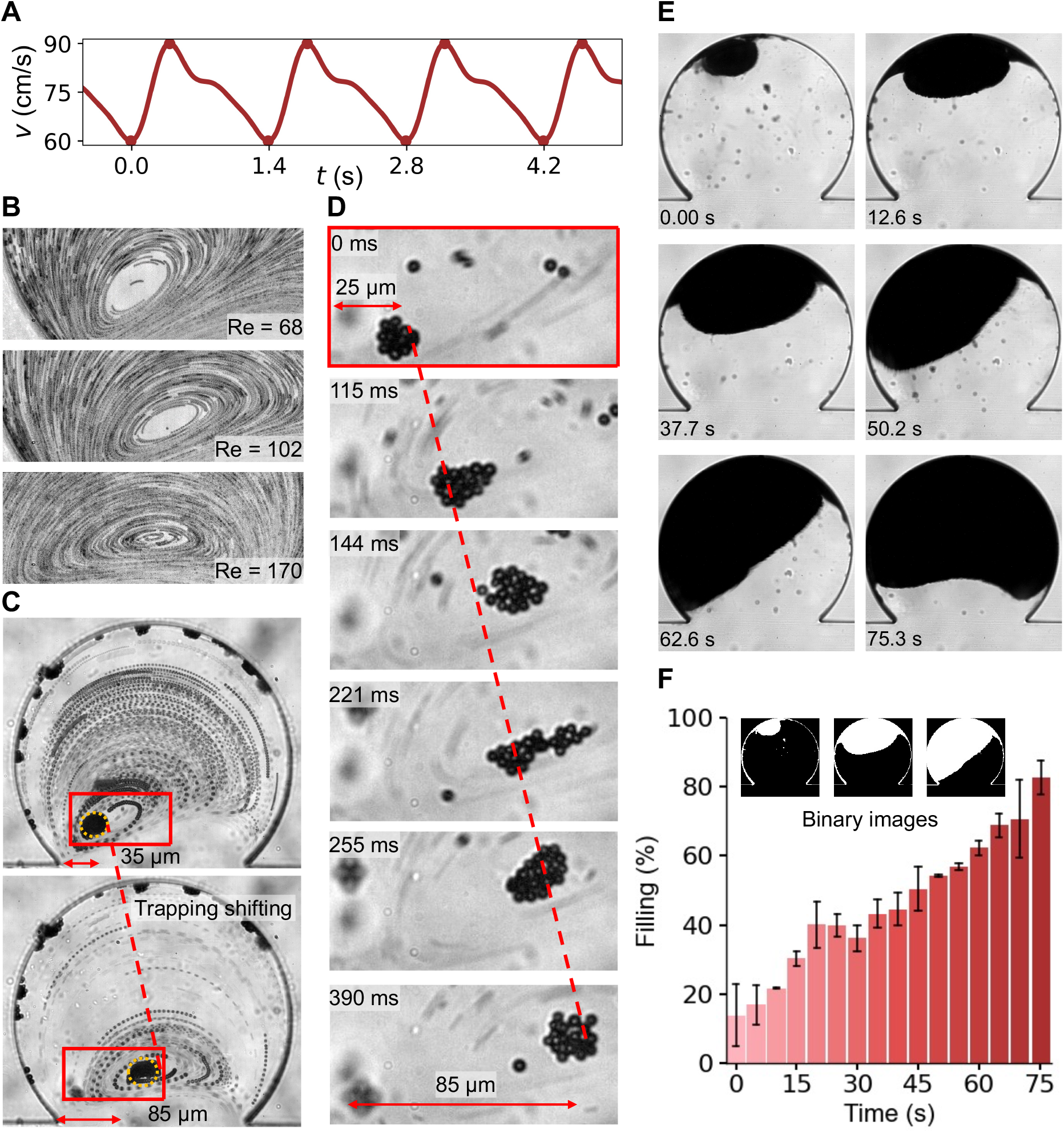
US-guided microbubble trapping and progressive cavity filling under disturbed flow. **(A)** Pulsatile flow profile showing a sinusoidal waveform at 60 bpm, with velocities oscillating between 60 and 80 cm s^−1^. **(B)** Experimental visualization of vortex core positions at different *Re*, illustrating shifts in the core location within the aneurysm cavity. **(C)** MB behavior under disturbed pulsatile flow with 1.7 MHz and 15 V_pp_ US activation, showing clusters trapped at the vortex eye and following the displacement of the vortex core. **(D)** Time-lapse sequence showing MB clusters entrained within the vortex and translating with the shifting vortex core under pulsatile flow. **(E)** Progressive accumulation of MBs within the aneurysm cavity under continuous US(2 MHz and 20 V_pp_) during pulsatile flow, visualized as a time-lapse sequence. **(F)** Quantification of cavity filling under pulsatile flow, represented as the fraction of white pixels within the aneurysm region from binarized image sequences.

## Discussion

Disturbed vascular flow is typically viewed as a pathological hallmark, associated with vortices that promote endothelial dysfunction, aneurysm growth, and plaque formation. Yet, our results show that these same disturbed flow fields, when coupled with US, can be harnessed to exert selective control over microscopic agents. We uncovered a previously unrecognized mechanism in which, under US activation, compressible MBs are drawn into vortex cores, undergo collective interactions, and are progressively deposited along cavity walls, whereas rigid particles and red blood cells bypass the trap. This establishes US–vortex coupling as a powerful framework for selective particle manipulation in complex hemodynamic environments.

Mechanistically, the observed behavior arises from the coupling between US and the disturbed flow field. In the absence of US, MBs follow the main streamlines and bypass the cavity. When US is applied, the acoustic radiation force directs bubbles from the channel into the cavity and biases their motion toward the low-pressure eye of the vortex. Within this region, secondary Bjerknes interactions between oscillating bubbles promote clustering and collective growth. As clusters enlarge, the balance between acoustic and viscous forces shifts until they are expelled from the vortex core and move toward the wall. Near the wall, image-mediated acoustic attraction promotes adhesion. Repetition of these capture, growth, ejection, and adhesion cycles results in progressive surface coverage of the cavity, a phenomenon that emerges only under the combined action of US and disturbed flow.

The clinical relevance of this mechanism is underscored by diseases in which disturbed flow is central (e.g., intracranial aneurysms^26,27^). Contemporary management remains largely catheter-based^28–31^ (clipping/coiling^26,32^, flow diversion^26^, liquid embolic injection^33,34^) with nontrivial procedural risks. Our results indicate that US can achieve stable wall engagement non-invasively by guiding compressible MBs into disturbed-flow regions and depositing them selectively on the cavity wall. Related work points to the importance of wall contact: pharmacological approaches^35^ such as cycloastragenol reduced aneurysm progression and preserved elastin in preclinical models^36^. More broadly, flow structures can act as selectors or triggers for delivery. In stenoses, shear-activated carriers disassemble in situ^37^, showing that hemodynamics itself can target therapy. By analogy, in aneurysm-like cavities, US leverages the recirculating vortex to transport and concentrate compressible MBs, enabling controlled deposition that fills the cavity. This principle can be extended: MBs with adhesive or polymerizable shells^35,38,39^ could convert transient attachment into durable filling without catheters; alternatively, focused US can cross-link deposited material in vivo (“sound printing”)^40^, providing a route to stabilize coverage after placement.

More broadly, this work establishes a framework for investigating and exploiting particle behavior in disturbed flows, a regime that has been difficult to access experimentally. By coupling US with recirculating vortices, we demonstrate selective trapping, clustering, and deposition that may extend beyond aneurysm-like cavities to drug delivery, clot modulation, or diagnostic imaging. Some limitations remain: our models are quasi-two-dimensional and employ simplified geometries, and translation to physiological conditions will require accounting for three-dimensional vascular architectures, variable flow patterns, and realistic boundary conditions. Future studies using 3D-printed vascular phantoms, animal models, and clinically compatible US systems will be essential to advance this approach toward in-vivo relevance.

## Code Availability

All data processing and analysis were performed using custom Python scripts. The core theoretical-model implementation (with supporting post-processing code) will be made publicly available on GitHub upon publication.

## Contributions

M.M. and D.A. conceived the project. D.A. supervised the project. M.M. discovered microbubbles trapping at the vortex eye. M.M. performed all the experiments and performed data analysis with feedback from N.N and D.A. N.N., M.M., and D.A contributed to the theoretical understanding and developed numerical simulations. All authors contributed to the experimental design, scientific presentation, discussion, and wrote the manuscript.

## Acknowledgments

This project has received funding from the European Research Council under the European Union’s Horizon 2020 Research and Innovation programme (Grant Agreement No. 853309, SONOBOTS), the Swiss NSF (Project Funding MINT 2022, Grant Agreement No. 213058) and an ETH Research Grant (Grant Agreement No. ETH-08 20-1). N.N acknowledges funding support from the United States National Science Foundation (OIA-2229636, CBET-2407937).

## References

1. Meng, H. et al. Complex Hemodynamics at the Apex of an Arterial Bifurcation Induces Vascular Remodeling Resembling Cerebral Aneurysm Initiation. Stroke 38, 1924–1931 (2007).

2. Chiu, J.-J. & Chien, S. Effects of Disturbed Flow on Vascular Endothelium: Pathophysiological Basis and Clinical Perspectives. Physiol Rev 91, 10.1152/physrev.00047.2009 (2011).

3. Ku, D. N. BLOOD FLOW IN ARTERIES. Annual Review of Fluid Mechanics 29, 399–434 (1997).

4. Taylor, C. A. & Draney, M. T. EXPERIMENTAL AND COMPUTATIONAL METHODS IN CARDIOVASCULAR FLUID MECHANICS. Annual Review of Fluid Mechanics 36, 197–231 (2004).

5. Arthurs, C. J. et al. CRIMSON: An open-source software framework for cardiovascular integrated modelling and simulation. PLOS Computational Biology 17, e1008881 (2021).

6. O’Reilly, M. A. Exploiting the mechanical effects of ultrasound for noninvasive therapy. Science 385, eadp7206 (2024).

7. Yan, J. et al. Reconfiguring active particles by electrostatic imbalance. Nature Mater 15, 1095–1099 (2016).

8. Li, J., Esteban-Fernández de Ávila, B., Gao, W., Zhang, L. & Wang, J. Micro/nanorobots for biomedicine: Delivery, surgery, sensing, and detoxification. Science Robotics 2, eaam6431 (2017).

9. Medany, M. M-Medany/Model-Based-Reinforcement-Learning-for-Ultrasound-Driven-Autonomous-Microrobots: Model-Based RL for US Microrobots – v1.0.0. Zenodo 10.5281/zenodo.15054076 (2025).

10. Real-time color flow mapping of ultrasound microrobots | Science Advances. https://www.science.org/doi/full/10.1126/sciadv.adt8887.

11. Shi, Z. et al. Ultrasound-Driven Programmable Artificial Muscles. bioRxiv 2024–01 (2024).

12. Di Carlo, D., Irimia, D., Tompkins, R. G. & Toner, M. Continuous inertial focusing, ordering, and separation of particles in microchannels. Proceedings of the National Academy of Sciences 104, 18892–18897 (2007).

13. Di Carlo, D., Edd, J. F., Humphry, K. J., Stone, H. A. & Toner, M. Particle Segregation and Dynamics in Confined Flows. Phys Rev Lett 102, 094503 (2009).

14. Carlo, D. D. Inertial microfluidics. Lab Chip 9, 3038–3046 (2009).

15. Vigolo, D., Radl, S. & Stone, H. A. Unexpected trapping of particles at a T junction. Proceedings of the National Academy of Sciences 111, 4770–4775 (2014).

16. Toner, M. & Irimia, D. Blood-on-a-chip. Annu Rev Biomed Eng 7, 77–103 (2005).

17. Ozkumur, E. et al. Inertial Focusing for Tumor Antigen–Dependent and –Independent Sorting of Rare Circulating Tumor Cells. Science Translational Medicine 5, 179ra47–179ra47 (2013).

18. Friend, J. Microscale acoustofluidics: Microfluidics driven via acoustics and ultrasonics. Rev. Mod. Phys. 83, 647–704 (2011).

19. Lenshof, A., Magnusson, C. & Laurell, T. Acoustofluidics 8: Applications of acoustophoresis in continuous flow microsystems. Lab Chip 12, 1210–1223 (2012).

20. Bruus, H. Acoustofluidics 7: The acoustic radiation force on small particles. Lab Chip 12, 1014–1021 (2012).

21. Sirsi, S. & Borden, M. Microbubble Compositions, Properties and Biomedical Applications. Bubble Sci Eng Technol 1, 3–17 (2009).

22. Crum, L. A. Bjerknes forces on bubbles in a stationary sound field. The Journal of the Acoustical Society of America 57, 1363–1370 (1975).

23. Leighton, T. G., Walton, A. J. & Pickworth, M. J. W. Primary Bjerknes forces. Eur. J. Phys. 11, 47 (1990).

24. Doinikov, A. A. Acoustic radiation forces: Classical theory and recent advances. Recent Res. Dev. Acoust 1, 39–67 (2003).

25. Pelekasis, N. A., Gaki, A., Doinikov, A. & Tsamopoulos, J. A. Secondary Bjerknes forces between two bubbles and the phenomenon of acoustic streamers. Journal of Fluid Mechanics 500, 313–347 (2004).

26. Toth, G. & Cerejo, R. Intracranial aneurysms: Review of current science and management. Vasc Med 23, 276–288 (2018).

27. Brisman, J. L., Song, J. K. & Newell, D. W. Cerebral Aneurysms. New England Journal of Medicine 355, 928–939 (2006).

28. Konda, R., Brumfiel, T. A., Bercu, Z. L., Grossberg, J. A. & Desai, J. P. Robotically steerable guidewires—Current trends and future directions. Science Robotics 10, eadt7461 (2025).

29. Rivkin, B. et al. Electronically integrated microcatheters based on self-assembling polymer films. Science Advances 7, eabl5408 (2021).

30. Gopesh, T. et al. Soft robotic steerable microcatheter for the endovascular treatment of cerebral disorders. Science Robotics 6, eabf0601 (2021).

31. Kim, Y. et al. Telerobotic neurovascular interventions with magnetic manipulation. Science Robotics 7, eabg9907 (2022).

32. Molyneux, A. J., Birks, J., Clarke, A., Sneade, M. & Kerr, R. S. C. The durability of endovascular coiling versus neurosurgical clipping of ruptured cerebral aneurysms: 18 year follow-up of the UK cohort of the International Subarachnoid Aneurysm Trial (ISAT). The Lancet 385, 691–697 (2015).

33. Panagiotopoulos, V. et al. Embolization of Intracranial Arteriovenous Malformations with Ethylene-Vinyl Alcohol Copolymer (Onyx). American Journal of Neuroradiology 30, 99–106 (2009).

34. Jin, D. et al. Swarming self-adhesive microgels enabled aneurysm on-demand embolization in physiological blood flow. Science Advances 9, eadf9278 (2023).

35. Shamloo, A., Ebrahimi, S., Amani, A. & Fallah, F. Targeted Drug Delivery of Microbubble to Arrest Abdominal Aortic Aneurysm Development: A Simulation Study Towards Optimized Microbubble Design. Sci Rep 10, 5393 (2020).

36. Wang, Y. et al. Inhibitory effects of cycloastragenol on abdominal aortic aneurysm and its related mechanisms. British Journal of Pharmacology 176, 282–296 (2019).

37. Shear-Activated Nanotherapeutics for Drug Targeting to Obstructed Blood Vessels | Science. https://www.science.org/doi/10.1126/science.1217815.

38. Makuta, T., Tamakawa, Y. & Endo, J. Hollow microspheres fabricated from instant adhesive. Materials Letters 65, 3415–3417 (2011).

39. Ferrante, E. A., Pickard, J. E., Rychak, J., Klibanov, A. & Ley, K. Dual targeting improves microbubble contrast agent adhesion to VCAM-1 and P-selectin under flow. J Control Release 140, 100–107 (2009).

40. Davoodi, E. et al. Imaging-guided deep tissue in vivo sound printing. Science 388, 616–623 (2025).

