## Supporting Information for "Ultrasound-induced Particle Dynamics in Pathological Vascular Vortices"

Mahmoud Medany , Nitesh Nama, and Daniel Ahmed*
[1]Acoustic Robotics Systems Lab, Institute of Robotics and Intelligent Systems, Department of Mechanical and Process Engineering, ETH Zurich, Switzerland
[2]Department of Mechanical & Materials Engineering, University of Nebraska-Lincoln, USA

### Material and Methods

Experimental setup

The microfluidic channels used in the study were produced through standard soft lithography using polydimethylsiloxane (PDMS). Each device was fabricated using a master mold lithographically patterned with SU-8 negative photoresist on a 4-inch silicon wafer, which was later placed inside a Petri dish. The thermocurable PDMS prepolymer was prepared by mixing the curing agent with the base at a weight ratio of 10:1. After degassing under vacuum, the prepolymer was cast onto the mold. PDMS was cross-linked by thermal curing for 2 h at 85 °C. The PDMS was poured into a mold and then cut and peeled from the channel mold. A 0.75 mm punch was used to form the inlet and outlet ports. The ports were created by mounting the puncher at a 60° angle, which prevented fluid from entering the channel at an orientation that could compromise the plasma-treatment bond between PDMS layers, thereby avoiding leakage and device malfunction. An additional PDMS layer was bonded to the PDMS channel by plasma treatment (1 min), followed by thermal annealing at 85 °C for 2 h. A piezo transducer was attached to the PDMS channel wall orthogonal to the aneurysm cavity. The channel flow was circulated using a controlled syringe pumps connected via silicone tubing to the channel inlet, and the effluent was collected in a 5 mL syringe. A Steminc piezoelectric ceramic plate was used as the US transducer. The actuator was bonded adjacent to the microfluidic chamber with a two-component epoxy to ensure efficient coupling to the PDMS wall, while avoiding any contact with the glass substrate. To mitigate acoustic impedance mismatch at the PDMS–air interface and suppress standing-wave formation, the entire device was submerged in a water bath. In addition to a baseline aneurysm cavity, five aneurysm geometries were implemented to span clinically relevant cases. The transducer was driven by a Tektronix arbitrary function generator (AFG3011C). The setup was mounted on an inverted microscope (Zeiss Axiovert 200m), and images were acquired with an AxioCAM MrM, a Photometrics HQ2 high-sensitivity camera, or a Chronos high-speed camera.

Microbubbles (MBs)

Microbubbles (MBs) were obtained from a clinical ultrasound contrast agent (SonoVue®, Bracco)(*1*). The kit includes a glass vial with 25 mg of lyophilized sulfur hexafluoride ($SF_6$) lipid-type A formulation, a prefilled 5 mL sterile saline syringe, and a Mini-Spike transfer set. For activation, saline was transferred into the vial using the supplied spike, followed by gentle agitation to yield a uniform dispersion of phospholipid-shelled MBs with $SF_6$ cores. The resulting population spans approximately 2–9 μm in diameter. Working suspensions at defined concentrations were prepared by diluting the activated agent with the provided saline to enable systematic testing across MB number densities.

Microparticle Tracers

Polystyrene microspheres (Polybead® Microspheres, Polysciences) with nominal diameters of 1.06 μm and 2.0 μm were used as flow tracers. The microspheres were suspended in saline by adding two drops of the stock particle solution to 5 mL of saline, corresponding to approximately 2.7% (v/v). The mixture was gently vortexed and briefly sonicated to ensure homogeneous dispersion and to prevent aggregation. The suspension was kept at room temperature (~25 °C) prior to use in flow characterization experiments.

Blood Samples

Mouse blood was used to evaluate microswarm behavior in hematological environments. Freshly drawn blood was anticoagulated with heparin and stored on ice until experiments were performed. In some trials, whole blood was introduced directly into the microfluidic channel; in others, the blood was diluted with sterile saline at volumetric ratios of 1:1 or 1:3 (blood:saline) to adjust viscosity and optical transparency. Where applicable, SonoVue® microbubbles were mixed with the blood or blood–saline mixture immediately before introduction to achieve the desired MB concentration.

Numerical Simulation

We characterized flow in a microfluidic aneurysm analogue using water at room temperature ($\rho \approx 997\,\mathrm{kg\,m^{-3}}$, $\mu \approx 0.89\,\mathrm{mPa{\cdot}s}$) driven by a syringe/peristaltic pump to impose defined volumetric flow rates. The device comprises a 50 μm-high parent channel connected to a 300 μm-diameter cavity and an 11 μm secondary channel. Flow was visualized with 2 μm tracer particles; their small response time (Stokes number $\ll 1$) ensures faithful tracking of the fluid, and time-averaged micrographs report the mean motion. Inlet bulk velocities of 5, 60, 107, and 120 $\mathrm{cm\,s^{-1}}$ correspond to Reynolds numbers $\mathrm{Re} = \rho \bar{u} D_h / \mu$of 5.5, 66, 117, and 132, where $D_h$ is the hydraulic diameter of the parent channel. Numerical simulations (COMSOL Multiphysics) solved the steady incompressible Navier–Stokes equations with water properties. Boundary conditions were a uniform normal inflow equal to the prescribed bulk velocity, no-slip walls, and zero-gauge outlet pressure; a fine mesh with boundary-layer refinement was used. The model reproduces the experiments across the Re range: a recirculation vortex forms in the cavity and strengthens with increasing Re, with the vortex core shifting toward the neck. Because the pump fixes flow rate, the parent-channel profile develops downstream and the local centerline speed reaches $\approx 2\,\bar{u}$ (Poiseuille effect), while the controlled quantities remain $\bar{u}$and Re. Streamline patterns and velocity-magnitude maps (color scale in $\mathrm{cm\,s^{-1}}$) thus provide a consistent basis for comparing conditions (Fig. S1)

Particle Image Velocimetry (PIV)

Flow in the microfluidic aneurysm analogue was quantified by cross-correlation PIV. Water at room temperature (~25 °C; $\rho \approx 997\,\mathrm{kg\,m^{-3}}$, $\mu \approx 0.89\,\mathrm{mPa{\cdot}s}$) was seeded with polystyrene microspheres (Polybead®, Polysciences; 1.06 μm and 2.0 μm), for which the Stokes number is $\ll 1$, ensuring faithful flow tracking. High-speed image sequences were recorded at 1100 fps on an inverted microscope under bright-field illumination. Velocity fields were computed in PIVlab (MATLAB) using $32 \times 32$ px interrogation windows with 50% overlap and sub-pixel Gaussian peak fitting. Spurious vectors were rejected by peak-ratio and local-median filters; validated fields were ensemble-averaged over the acquisition window, and vorticity and streamlines were derived from the final vector maps.

**Supporting Text 1**
**Model for capture, clustering, and ejection under ultrasound**

In continuous flow, gas-filled microbubbles (MBs; 2.5 μm) bypass the aneurysm cavity without ultrasound. With ultrasound (US) on, MBs are redirected into the cavity and, once inside, follow elliptical paths that spiral into the vortex eye; the speed decreases from ~3 mm $s^{-1}$ near the entrance to ~0.5 mm $s^{-1}$ close to the eye, culminating in stable trapping (Fig. 2A–C). A minimal, data-driven model reproduces this guidance and explains the subsequent clustering and ejection.

Governing balance (overdamped slip)

For a bubble or cluster of effective radius $R_c$ and volume $V_c = \frac{4}{3}\pi R_c^3$ at position $\mathbf{x}(t)$with velocity $\mathbf{v}(t)$, the force balance is

$$6\pi\mu R_c(\mathbf{v} - \mathbf{u}_f(\mathbf{x})) = \; -V_c \nabla p_{\text{vort}}(\mathbf{x}) + \mathbf{F}_{rad} + \mathbf{F}_B. \qquad \text{(S1)}$$

The primary radiation force is summarized as

$$\mathbf{F}_{rad}(t) \approx \mathcal{G}_{ac}\, V_c(t)\, \hat{\mathbf{g}}, \qquad \text{(S2)}$$

where $\mu$ is the dynamic viscosity; $\mathbf{u}_f(\mathbf{x})$ is the measured carrier flow (COMSOL); $p_{\text{vort}}(\mathbf{x})$ is a Rankine-type pressure surrogate with a minimum at the eye; $\hat{\mathbf{g}}$ points from the transducer toward the opposite wall; and $\mathcal{G}_{ac}$ (Pa $m^{-1}$) bundles the effective acoustic contrast with the mean pressure-squared gradient. Because $\mathbf{F}_{rad} \propto V_c(t)$, the acoustic bias strengthens as clusters grow. The term $\mathbf{F}_B$ captures secondary Bjerknes attractions among oscillating MBs in the core and image-Bjerknes(*2*) pull near a wall.

Vortex confinement (trap)

Let the eye be at $\mathbf{x}_c = (x_c, y_c)$, with core radius $a$(radius of peak speed) and circulation $\Gamma$; define $\Omega = \Gamma/(2\pi a^2)$ and $r = \parallel \mathbf{x} - \mathbf{x}_c \parallel$. A Rankine surrogate for the outward radial pressure gradient is

$$\partial_r p_{\text{vort}}(r) = \begin{cases} \rho\, \Omega^2\, r, & r < a, \\ \rho\, \Gamma^2/(4\pi^2 r^3), & r \geq a, \end{cases} \qquad \text{(S3)}$$

so $p_{\text{vort}}$ has a minimum at the eye and the confinement term $\mathbf{F}_v = -V_c \nabla p_{\text{vort}}$ points inward, cooperating with $\mathbf{F}_{rad}$ to draw MBs into the core.

Using the measured $\mathbf{u}_f$ and Eqs. (S1)–(S3), releasing MBs from distinct entry positions reproduces inward spiraling and convergence to the eye (Fig. 2D). The full field with simulated tracks shows capture is robust across entry angles (Fig. 2E).

Selectivity (MBs vs. rigid particles/RBCs)

In controls, polystyrene tracers (1 μm) and RBCs follow streamlines and do not trap under US, whereas MBs are redirected and confined to the eye (Fig. 3A–D)(Fig. S4) Mechanistically, gas-filled MBs are highly compressible and oscillate under US, experiencing strong primary radiation and secondary-Bjerknes attractions(*2*) that bias them toward the low-pressure eye; rigid particles and RBCs are comparatively incompressible, respond weakly to US, and are governed mainly by inertial drag past the eye.

General ejection criterion (projected balance)

Because the US-driven acoustic forces on an MB cluster—the primary radiation force and the secondary/image-Bjerknes(*2*) term—grow approximately with cluster volume ($\mathbf{F}_{rad} \sim V_c \propto R_c^3$) (*3–5*), whereas the viscous resistance grows linearly with size ($F_D = 6\pi\mu R_c\, U$), larger clusters experience an increasing outward slip relative to the background recirculation. Ejection occurs once the net acoustic drive exceeds hydrodynamic confinement and drag:

$$\| \mathbf{F}_{rad} + \mathbf{F}_B \| \ \gtrsim\ V_c\ \| \nabla p_{\text{vort}} \| \ +\ 6\pi\mu R_c\, U. \qquad \text{(S4)}$$

Why confinement is negligible near the eye

In the Rankine core, $\partial_r p_{\text{vort}} = \rho\, \Omega^2 r$is linear in $r$and vanishes at the eye. Consequently,

$$\frac{| \mathbf{F}_v |}{| \mathbf{F}_{rad} |} = \frac{V_c\ | \nabla p_{\text{vort}} |}{\mathcal{G}_{ac}\, V_c} = \frac{\rho\, \Omega^2\, r}{\mathcal{G}_{ac}}\ \ll\ 1 \text{for}\ r \ll a, \qquad \text{(S5)}$$

so near the eye the balance is dominated by acoustic drive versus Stokes drag. Under our fixed drive, this condition is met at a reproducible critical size ($\sim$ 15 $\mu$m; Fig. 3E–F), after which clusters depart the eye and traverse the cavity.

Clustering, growth, and ejection sequence

Once several MBs reach the eye, ultrasound-driven interactions dominate: secondary-Bjerknes attraction and image-mediated wall attraction(*2*) promote hierarchical assembly (doublets → triplets → larger aggregates) within milliseconds (Fig. 3A–C; Fig. S3). Clusters grow at ~6 MB $s^{-1}$ (≈ 30 $\mu m^2\ s^{-1}$ area increase) and, under fixed drive (1.04 MHz, 18.1 Vpp), reach a critical radius ∼15 µm, beyond which the trap destabilizes and the aggregate is abruptly ejected (Fig. 3D). Ejected clusters traverse the cavity and anchor at the opposite wall via image-bubble attraction; RBCs continue to bypass the trap. Ejection speeds begin around ~0.25 mm $s^{-1}$ and rise to ∼1 mm $s^{-1}$ just before reaching the aneurysm wall (Fig. S5). Quantification shows a consistent size threshold (Fig. 3E–F).

Connection to Gor'kov (meaning of $\mathcal{G}_{ac}$)

Writing the primary radiation force as $\mathbf{F}_{rad} = -\nabla U_G$ shows $\mathcal{G}_{ac} = K_{ac}\ \| \nabla\langle p^2 \rangle \|$, where $K_{ac} = (f_1 - \frac{3}{2} f_2)/(2\rho_0 c_0^2)$and $f_{1,2}$ are effective (shell- and frequency-dependent) contrast factors. We do not map $\| \nabla\langle p^2 \rangle \|$directly here; $\mathcal{G}_{ac}$ is used as a lumped parameter in Eqs. (S1)–(S5).

**Supporting Text 2**
**Numerical solver and code implementation**

We simulate the capture stage of two independent microbubbles (MBs) released into the measured 2-D carrier flow of the aneurysm model. The model uses a Rankine-type vortex pressure surrogate with a minimum at the eye and a slip-to-flow drag that rapidly relaxes the MB velocity toward the local carrier flow. Bubble–bubble forces (secondary Bjerknes(*2*)), wall images, and explicit primary radiation forcing are intentionally omitted here; those are treated in the clustering/ejection analysis. This solver reproduces the spiral-in and trapping across entry angles (Fig. 2D–E).

For each bubble $i = 1,2$ with position $\mathbf{x}_i = (x_i, y_i)$and velocity $\mathbf{v}_i = (u_i, v_i)$:

$$\dot{\mathbf{x}}_i = \mathbf{v}_i, \dot{\mathbf{v}}_i = -\frac{3}{\rho} \nabla p_{\text{rank}}(\mathbf{x}_i; \mathbf{x}_c, a, \Gamma) \; + \; 0.75\, C_D\, (\mathbf{u}_f(\mathbf{x}_i) - \mathbf{v}_i) \parallel \mathbf{u}_f(\mathbf{x}_i) - \mathbf{v}_i \parallel.$$

Here $\mathbf{u}_f$is the measured carrier field from CSV (interpolated by k-nearest neighbors), $p_{\text{rank}}$is a Rankine surrogate, $C_D$is an effective drag constant, and $\rho$is the fluid density. Important: keep the coefficients exactly as above (this matches the code).
Rankine pressure surrogate**.** With $r = \parallel \mathbf{x} - \mathbf{x}_c \parallel$and $\Omega = \Gamma/(2\pi a^2)$,

$$\partial_r p_{\text{rank}}(r) = \begin{cases} \rho\, \Omega^2\, r = \dfrac{\rho\, \Gamma^2}{4\pi^2 a^4}\, r, & r < a, \\ \dfrac{\rho\, \Gamma^2}{4\pi^2 r^3}, & r \geq a, \end{cases} \qquad \nabla p_{\text{rank}} = \partial_r p_{\text{rank}}\, \hat{\mathbf{r}}.$$

<u>Time integration and events.</u>
ODEs were solved in Python with *scipy.integrate.solve_ivp* (*6*, *7*) using the explicit RK45 method(*8*) with tolerances rtol = $1\times10^{-6}$ and atol = $1\times10^{-9}$. An event terminated the integration if a bubble left the axis-aligned bounding box of the measured field. For Windows spawn-safety and to bound step sizes, the full horizon was partitioned into $N$contiguous subintervals ($N$ = CPU count) and integrated sequentially; each subinterval started from the previous subinterval's terminal state, preserving trajectory continuity.

---

Algorithm 1: Two-microbubble trajectory solver

**Require:** CSV $(x, y, u, v)$, parameters $\rho, C\_D, \Gamma$, a; initial positions $(x_0, y_0)$; tolerances $rtol, atol$
**Ensure:** Trajectory file trajectory_two_mb.csv

Read velocity CSV → build cKD Tree (points) for k-NN interpolation of $\mathbf{u}_f(x, y)$.
Estimate vortex center $(x\_c, y\_c)$ by minimizing radial velocity component.
Set core radius a from peak-speed radius or $a\_override$.
Define Rankine pressure gradient:
  $\partial r\; p \;=\; (\rho\Gamma^2/4\pi^2)(r/a^4)\;\; if\;\; r < a,$
  $(\rho\Gamma^2/4\pi^2)(1/r^3)\;\; if\;\; r \geq a.$
Initialize state $Y \;=\; [x1, y1, u1, v1, x2, y2, u2, v2]$.
For each time-chunk:
  integrate $\dot{Y} \;=\; f(t, Y)$ with $solve\_ivp$ (RK45, $rtol \;=\; 1e - 6, atol \;=\; 1e - 9$);
  stop if bubble leaves $[xmin, xmax] \times [ymin, ymax]$;
  seed next chunk with final state.
Concatenate results → save CSV of $(x\_i(t), y\_i(t), u\_i(t), v\_i(t))$.
Render quiver plots of measured field + simulated trajectories

---

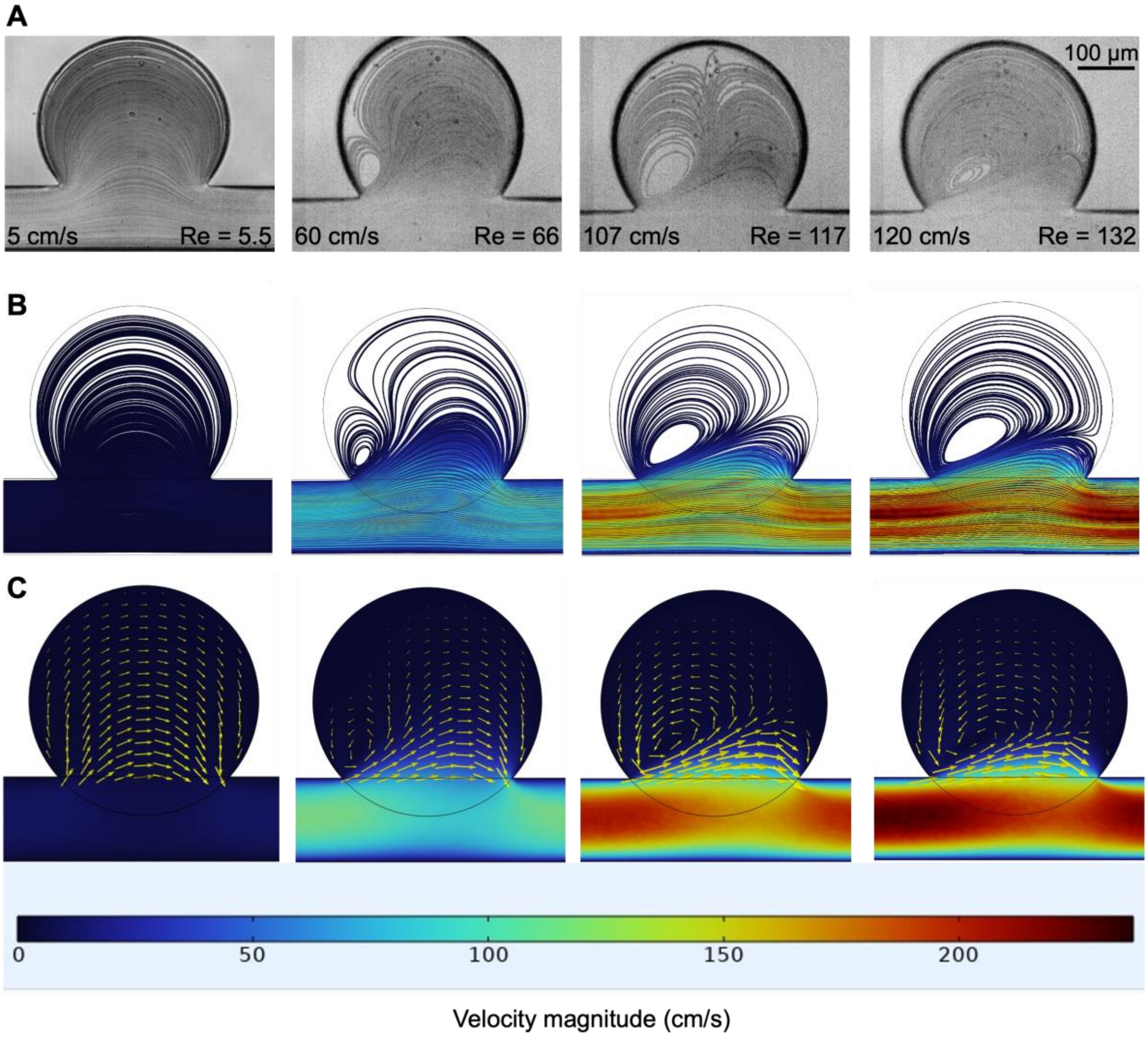


**Fig. S1. Transition from laminar to disturbed flow in a microaneurysm analogue. (A)** Time-averaged micrographs of 2 µm tracers at inlet bulk speeds 5, 60, 107, and 120 cm $s^{-1}$ (Re = 5.5, 66, 117, 132). Recirculation emerges and the vortex core shifts toward the neck as Re increases. Scale bar, 100 µm. **(B–C)** 2D-extruded COMSOL simulations of the same geometry (parent-channel height 50 µm, 300 µm cavity, 11 µm secondary channel) showing streamlines and velocity magnitude with vectors (cm $s^{-1}$). Boundary conditions: uniform normal inflow (bulk velocity), no-slip walls, zero-gauge outlet pressure. The developed Poiseuille profile produces centerline speeds ≈ 2× the bulk value and reproduces the vortex dynamics observed experimentally.

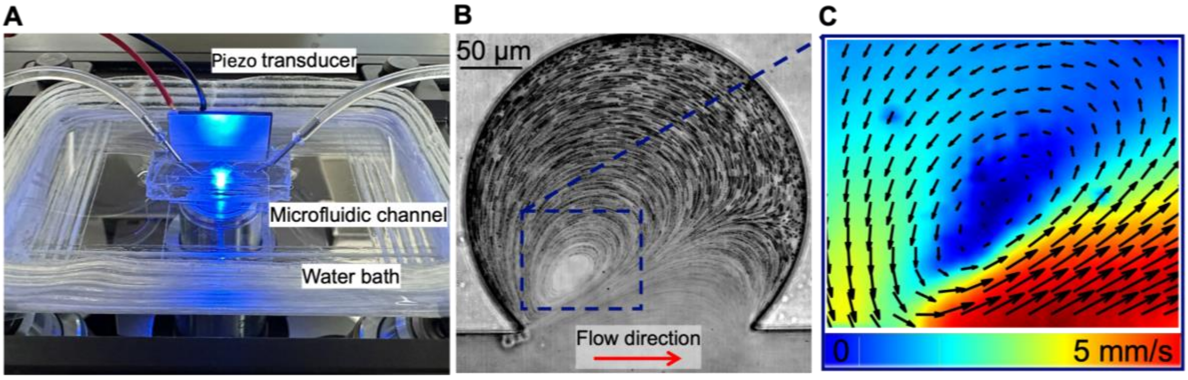


**Fig. S2. Experimental setup and PIV analysis of vortex flow in the aneurysm cavity. (A)** Microfluidic device mounted on an inverted microscope: the PDMS channel is submerged in a water bath for acoustic coupling, and a piezoelectric plate transducer is bonded to the channel wall. **(B)** Representative high-speed bright-field image of the aneurysm cavity seeded with polystyrene tracer particles (1.06 µm); flow left→right. Dashed box marks the region analyzed. **(C)** Time-averaged PIV field from a sequence recorded at 1100 fps (≈25 °C), showing in-plane velocity vectors overlaid on a velocity-magnitude colormap. The map reveals a recirculation vortex occupying the dome with a low-speed core and increasing shear toward the neck. PIV was performed in PIVlab (MATLAB) as described in Methods.

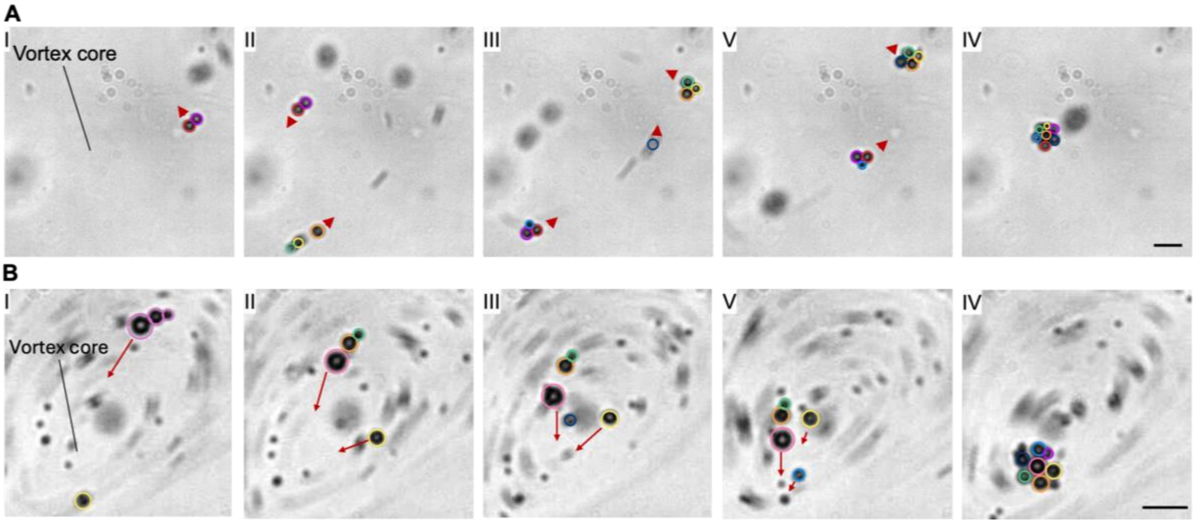


**Fig. S3. Cooperative microbubble assembly at the vortex core under ultrasound. Top row (I–V):** Time-lapse frames showing initially separate microbubbles (colored outlines) approaching the aneurysm vortex; small doublets/triplets form en route (red arrowheads) as bubbles enter the recirculation zone. Bottom row **(I–V)**: As the group nears the vortex core, inward spiraling trajectories (red arrows) bring the pre-assembled pairs/triads together, merging into a single compact cluster at the core. This example demonstrates an alternative pathway to that in Fig. 3: assembly can proceed by simultaneous arrival and fusion of small clusters, not only by stepwise monomer addition. Ultrasound was continuously on. Scale bar, 15 µm.

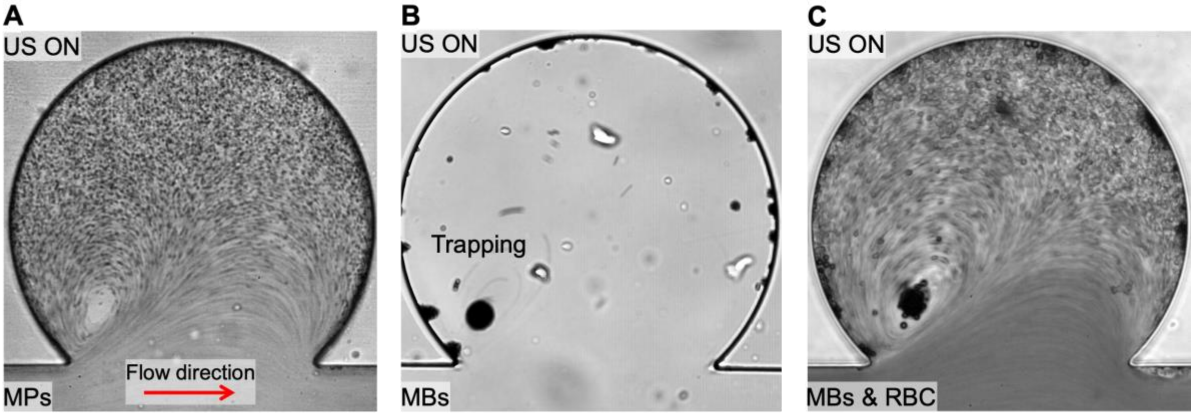


**Fig. S4. Selective ultrasonic trapping of microbubbles at the vortex core. (A)** Polystyrene microparticles (1.06 µm; MPs) with ultrasound **ON** circulate with the recirculating streamlines but do **not** localize at the vortex eye. **(B)** Under the same conditions, SonoVue microbubbles (MBs) concentrate and are **trapped** at the vortex core. **(C)** In a mouse blood–saline mixture, MBs again localize at the vortex eye while red blood cells (RBCs) continue to recirculate without trapping. All panels: flow from left to right. Scale bar, 50 µm.

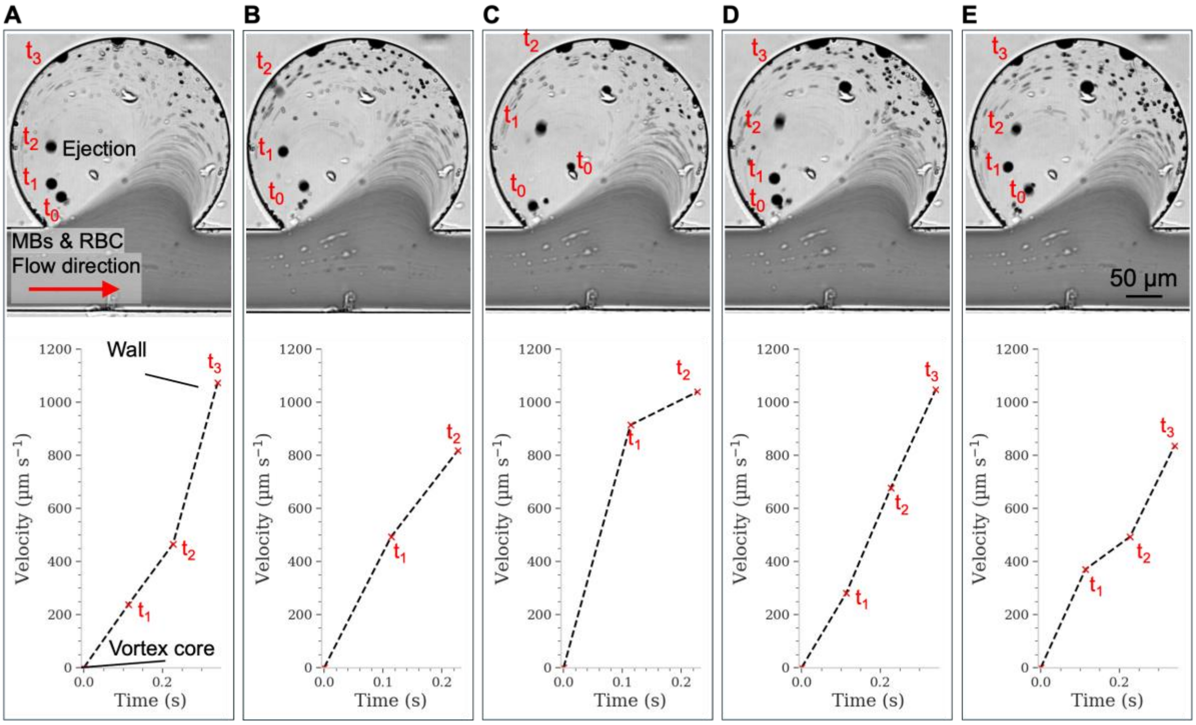


**Fig. S5. Ultrasound-driven ejection dynamics in a blood–saline environment (five repeats). (A–E)** Top: time-lapse micrographs of a mouse blood–saline mixture containing microbubbles in the aneurysm cavity under continuous ultrasound (1.6 MHz; US on). Flow enters left→right (arrow). The tracked aggregate (MBs) is marked at successive times $t_0$–$t_3$ as it is ejected from the vortex core toward the cavity wall. Bottom: corresponding speed–time traces for each repeat; points at $t_0$–$t_3$ match the image annotations and dashed lines are guides to the eye. Across n = 5 independent runs, ejection consistently occurs with an acceleration phase followed by rapid translation along the wall, with trial-to-trial variability in peak speed.

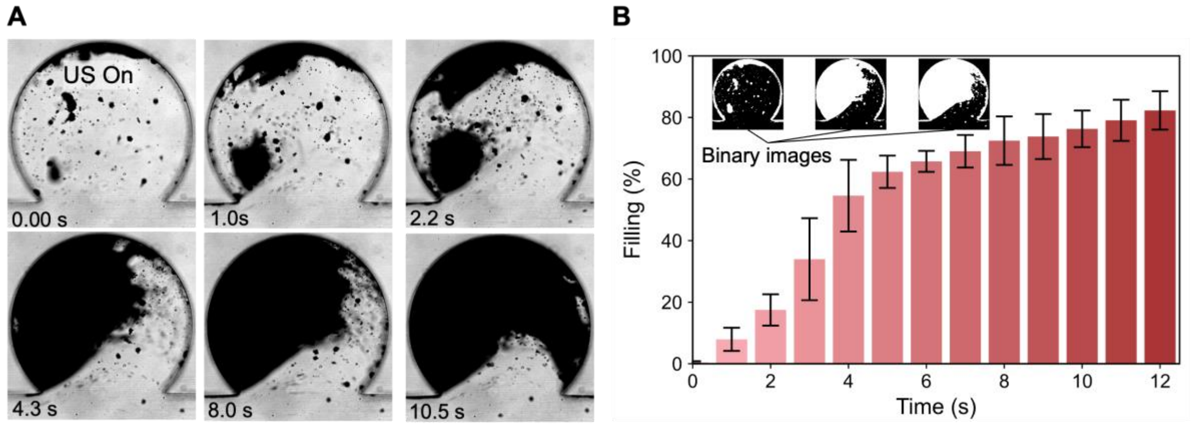


**Figure S6. Continuous ultrasound-driven filling of the aneurysm cavity. (A)** Sequential micrographs showing the continuous accumulation of microbubbles (4 µL mL$^{-1}$) within the aneurysm cavity at an inlet velocity of 60 cm s$^{-1}$ under continuous ultrasound exposure (2.03 MHz, 20 $V_{pp}$). Microbubbles progressively occupy the recirculation zone, resulting in monotonic filling of the dome over ≈ 10 s after ultrasound onset. **(B)** Quantification of cavity filling as a function of time, derived from binary segmentation of the image sequence (example binary images shown in the inset). The filling fraction increases steadily with exposure time, reaching nearly 90% cavity occupation after ≈ 12 s.

### References


1. F. Piscaglia, L. Bolondi, The safety of Sonovue® in abdominal applications: Retrospective analysis of 23188 investigations. *Ultrasound in Medicine & Biology* **32**, 1369–1375 (2006).
2. N. A. Pelekasis, A. Gaki, A. Doinikov, J. A. Tsamopoulos, Secondary Bjerknes forces between two bubbles and the phenomenon of acoustic streamers. *Journal of Fluid Mechanics* **500**, 313–347 (2004).
3. L. A. Crum, Bjerknes forces on bubbles in a stationary sound field. *The Journal of the Acoustical Society of America* **57**, 1363–1370 (1975).
4. T. G. Leighton, A. J. Walton, M. J. W. Pickworth, Primary Bjerknes forces. *Eur. J. Phys.* **11**, 47 (1990).
5. A. A. Doinikov, Acoustic radiation forces: Classical theory and recent advances. *Recent Res. Dev. Acoust* **1**, 39–67 (2003).
6. W. McKinney, Data Structures for Statistical Computing in Python. *SciPy 2010*, doi: 10.25080/Majora-92bf1922-00a (2010).
7. P. Virtanen, R. Gommers, T. E. Oliphant, M. Haberland, T. Reddy, D. Cournapeau, E. Burovski, P. Peterson, W. Weckesser, J. Bright, S. J. van der Walt, M. Brett, J. Wilson, K. J. Millman, N. Mayorov, A. R. J. Nelson, E. Jones, R. Kern, E. Larson, C. J. Carey, İ. Polat, Y. Feng, E. W. Moore, J. VanderPlas, D. Laxalde, J. Perktold, R. Cimrman, I. Henriksen, E. A. Quintero, C. R. Harris, A. M. Archibald, A. H. Ribeiro, F. Pedregosa, P. van Mulbregt, SciPy 1.0: fundamental algorithms for scientific computing in Python. *Nat Methods* **17**, 261–272 (2020).
8. J. D. Hunter, Matplotlib: A 2D Graphics Environment. *Computing in Science & Engineering* **9**, 90–95 (2007).
